# Data-Driven Identification of Sex Differences in Cerebral Blood Flow Using Arterial Spin Labelling and Explainable Artificial Intelligence

**DOI:** 10.64898/2026.07.05.736642

**Authors:** Ninad Aithal, Neelam Sinha, Venkatesh Babu Radhakrishnan

**Affiliations:** Indian Institute of Science, Bangalore

**Keywords:** Cerebral Blood Flow, Arterial Spin Labelling, Sex Differences, SHAP, Explainable AI, Brainnetome Atlas, Deep Learning

## Abstract

**Purpose:** To investigate sex differences in cerebral blood flow through densely parcellated cortical and subcortical regions using explainable artificial intelligence methods and identify neurobiologically interpretable perfusion biomarkers.

**Methods:** High-resolution pseudo-continuous arterial spin labelling (1.875 mm × 1.875 mm × 3 mm) and structural MRI data were curated from 215 healthy young adults (150 females, 95 males; age 18–30 years) from the publicly available *I See your Brains* (ISYB) dataset. Cerebral blood flow was quantified using atlas-based regional analysis with the Brainnetome Atlas (246 regions) and optimized registration procedures. Sex classification employed diverse machine learning paradigms including linear classifiers, ensemble methods, and kernel-based approaches for regional CBF features, with deep convolutional neural networks (CNN) applied to whole-brain 3D imaging data. Model interpretability was achieved using SHapley Additive exPlanations (SHAP), computed over an ensemble of 500 logistic regression models (100 iterations × 5-fold cross-validation). Regions appearing among the top 20% of discriminative features more than 289 times were considered statistically significant using binomial testing. GradCAM was used to obtain class-specific attribution maps from the CNN model.

**Results:** Perfusion-based features demonstrated superior sex classification performance compared to structural morphometry. Regional CBF analysis using logistic regression achieved 91 ± 2% balanced accuracy and 0.95 ± 0.05 ROC-AUC, substantially outperforming morphometric features (85 ± 8% balanced accuracy, 0.88 ±0.06 ROC-AUC). Deep learning classification of 3D CBF maps achieved a performance of 92 ±5% balanced accuracy, 0.92 ± 0.05 ROC-AUC. SHAP analysis identified 30 statistically significant aggregation-agnostic CBF-based biomarker regions using regional CBF, predominantly involving frontoparietal control networks (27%) and default mode networks (17%). Grad-CAM revealed that the 3D CNN model primarily focused on regions within the frontal lobe. Morphometry-based analysis identified 28 discriminative regions with markedly different anatomical distribution (*r* = 0.21) emphasizing visual (32%) and default mode (14%) networks.

**Conclusion:** Cerebral blood flow patterns provide highly sensitive and biologically interpretable markers of sex differences in young adult brain. The identification of robust perfusion biomarkers through explainable AI demonstrates the clinical potential of ASL imaging for precision medicine applications in neuroscience. We establish a methodological framework for investigating sexspecific brain physiology using non-invasive neuroimaging.

## 1 Introduction

Understanding biological sex differences in brain structure and function has emerged as a critical frontier in precision medicine and neuroscience research. Sex plays a fundamental role throughout the lifespan, influencing early brain development, adolescent maturation, and aging processes, with profound implications for both normative brain functioning and pathological conditions [1, 2, 3]. The neurobiological foundations of these sex differences undergo dynamic changes across development, with structural and functional brain differences remaining relatively modest during early childhood but becoming increasingly pronounced during adolescence – a critical period characterized by extensive neural remodelling and the emergence of many neuropsychiatric conditions [4, 5]. This developmental window represents a particularly important timeframe for investigating sex-specific brain characteristics, as it coincides with divergent physical, behavioural, and psychological trajectories between males and females that may establish lifelong patterns of brain function and disease vulnerability [6, 7].

Cerebral blood flow (CBF), quantified as the volume of blood delivered per 100 grams of brain tissue per minute, serves as a fundamental physiological parameter reflecting metabolic demand, vascular integrity, and neural activity. Normal CBF ranges from 45–60 mL/100g/min and is regulated through complex interactions involving autoregulation, cerebral perfusion pressure, temperature and blood viscosity [8]. Arterial spin labelling (ASL) magnetic resonance imaging has revolutionized CBF assessment by providing non-invasive, quantitative perfusion measurements using magnetically labelled arterial water as an endogenous tracer [9, 10, 11]. Despite these advantages, routine clinical implementation of ASL remains limited, and the field lacks comprehensive, data-driven approaches to identify sex-specific perfusion biomarkers and elucidate underlying neurobiological mechanisms. Previous ASL investigations have consistently documented significant sex differences in cerebral perfusion patterns. Numerous studies have demonstrated that females exhibit substantially higher CBF values than males across both global and regional measurements [12, 13, 14, 15, 16, 17, 18, 19, 20]. However, these studies employed relatively coarse regional parcellations and primarily relied on statistical testing approaches, with modest sample sizes limiting their ability to identify specific neuroanatomical substrates and develop robust predictive biomarkers.

Data-driven approaches have achieved 70–90% accuracy in neuroimaging-based sex classification using structural and functional features [21, 22, 23]. While numerous studies have documented sex differences in cerebral perfusion, datadriven classification approaches for perfusion imaging remain underexplored. Existing perfusion studies often reduce rich spatial CBF information to summary statistics, potentially missing subtle discriminative patterns that machine learning could identify as neurobiological biomarkers of sex differences [24]. The increasing sophistication of machine learning models necessitates robust interpretability frameworks to enhance scientific understanding beyond predictive accuracy. SHapley Additive exPlanations (SHAP) provides a principled approach for quantifying individual feature contributions to model predictions while ensuring local accuracy and consistency, yet its application to perfusion-based sex classification remains unexplored [25, 26].

The present study addresses these limitations through a comprehensive, data-driven investigation of sex differences in cerebral perfusion using high-resolution ASL imaging in young adults (18–30 years). We leverage fine-grained regional parcellation (246 regions from the Brainnetome Atlas) to examine CBF patterns with high anatomical specificity, employing data-driven approaches for sex classification. Our analytical framework integrates SHAP-based interpretability to identify neurobiologically meaningful perfusion biomarkers, while systematic comparison with structural neuroimaging data elucidates the relationship between morphometric and perfusion-based sex differences. Our approach (Figure 1) aims to advance understanding of sex-specific brain physiology during a critical developmental period and demonstrate the clinical utility of ASL perfusion imaging for precision medicine applications.

**Figure 1.**
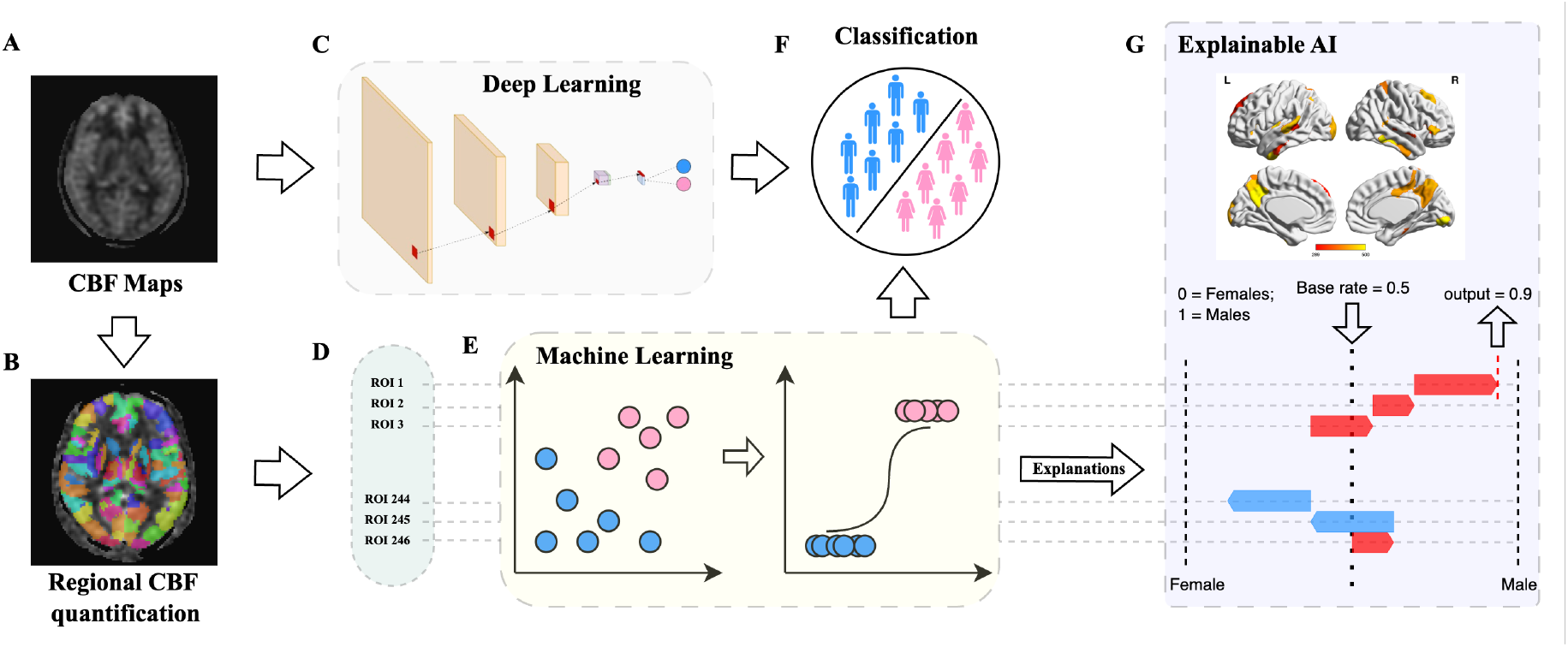
Study Overview. End-to-end framework for data-driven, explainable sex classification from cerebral blood flow.

## 2 Methods

### 2.1 Study Population and Imaging Protocol

We analyzed data from the publicly available “Imaging Chinese Young Brains” (ISYB) dataset, comprising 251 healthy Chinese Han participants from northern China (156 females, 95 males; age range: 18–30 years). Imaging quality control procedures are described in Gao et al. 2022 [27], and demographics are provided in Table 1.

**Table 1.** Participant demographics and brain morphometric characteristics. Stratified by biological sex; values are mean (SD). Comparisons via two-sample *t*-tests. S = significant (*p <* 0.05); NS = not significant.

| Characteristic | Males ($n = 95$) | Females ($n = 150$) | Sig. |
| --- | --- | --- | --- |
| Age (years) | 22.72 (2.78) | 22.50 (2.60) | NS |
| Total intracranial volume (mm<sup>3</sup>) | $1.60 \times 10^6$ | $1.45 \times 10^6$ | S |
| Total brain volume (mm<sup>3</sup>) | $1.25 \times 10^6$ | $1.15 \times 10^6$ | S |
| Total gray matter (mm<sup>3</sup>) | $7.20 \times 10^5$ | $6.66 \times 10^5$ | S |
| Total white matter (mm<sup>3</sup>) | $4.77 \times 10^5$ | $4.36 \times 10^5$ | S |
| Cortical gray matter (mm<sup>3</sup>) | $5.40 \times 10^5$ | $5.00 \times 10^5$ | S |
| Brain stem (mm<sup>3</sup>) | $2.10 \times 10^4$ | $1.94 \times 10^4$ | S |
| Total hippocampus (mm<sup>3</sup>) | 8842.75 | 8434.10 | S |
| Total amygdala (mm<sup>3</sup>) | 3780.40 | 3466.03 | S |
| Total thalamus (mm<sup>3</sup>) | $1.66 \times 10^4$ | $1.57 \times 10^4$ | S |
| Total caudate (mm<sup>3</sup>) | 7890.16 | 7435.21 | S |
| Total putamen (mm<sup>3</sup>) | $1.14 \times 10^4$ | $1.07 \times 10^4$ | S |
| Total pallidum (mm<sup>3</sup>) | 4291.20 | 4023.88 | S |
| Total accumbens (mm<sup>3</sup>) | 1264.81 | 1265.19 | NS |
| Total cerebellum WM (mm<sup>3</sup>) | $3.10 \times 10^4$ | $3.01 \times 10^4$ | NS |
| Total cerebellum cortex (mm<sup>3</sup>) | $1.16 \times 10^5$ | $1.06 \times 10^5$ | S |
| Avg. cortical thickness (mm) | $2.62 \times 10^4$ | $2.39 \times 10^4$ | S |
| Total cortical surface area (mm<sup>2</sup>) | $6.04 \times 10^6$ | $5.51 \times 10^6$ | S |

All scans were acquired on a 3.0 Tesla GE MR750 scanner. High-resolution T1-weighted images were collected using a 3D spoiled gradient-recalled echo sequence with the following parameters: matrix = 256× 256, 188 slices, field of view = 256 × 256 mm^2^, TR = 8.16 ms, TE = 3.18 ms, flip angle = 12^*°*^, inversion time = 450 ms. Pseudo-continuous ASL (PCASL) images were acquired with: matrix = 128 ×128, 50 slices, field of view = 240 ×240 mm^2^, TR = 5046 ms, TE = 11.09 ms, flip angle = 111^*°*^, inversion time = 2025 ms. Structural images were evaluated using the MRIQC toolkit, and cases with motion artifacts or anomalies were excluded.

### 2.2 Image Preprocessing

T1-weighted images were processed using FreeSurfer v5.3 [28] for cortical reconstruction and subcortical segmentation. The Brainnetome Atlas was registered to individual reconstructions for cortical parcellation and subcortical segmentation, yielding 666 morphometric features per subject: surface area, cortical thickness, and volume for 210 cortical regions, and volume for 36 subcortical regions.

A multi-step atlas registration pipeline (Figure 2) was implemented for CBF images using FSL’s Oxford ASL Toolbox [29]. T1-weighted images were reoriented, bias-corrected, skull-stripped, and registered to corresponding ASL images to derive individual T1-to-ASL transforms. Simultaneously, each T1-weighted image was non-linearly aligned to MNI152 space, allowing inverse transformation of the Brainnetome Atlas to subject space and subsequent mapping to ASL space. This enabled voxel-wise regional correspondence between CBF maps and anatomical ROIs. Mean, median, and maximum CBF values were extracted from each of the 246 atlas regions, generating 246-dimensional feature vectors per subject.

**Figure 2.**
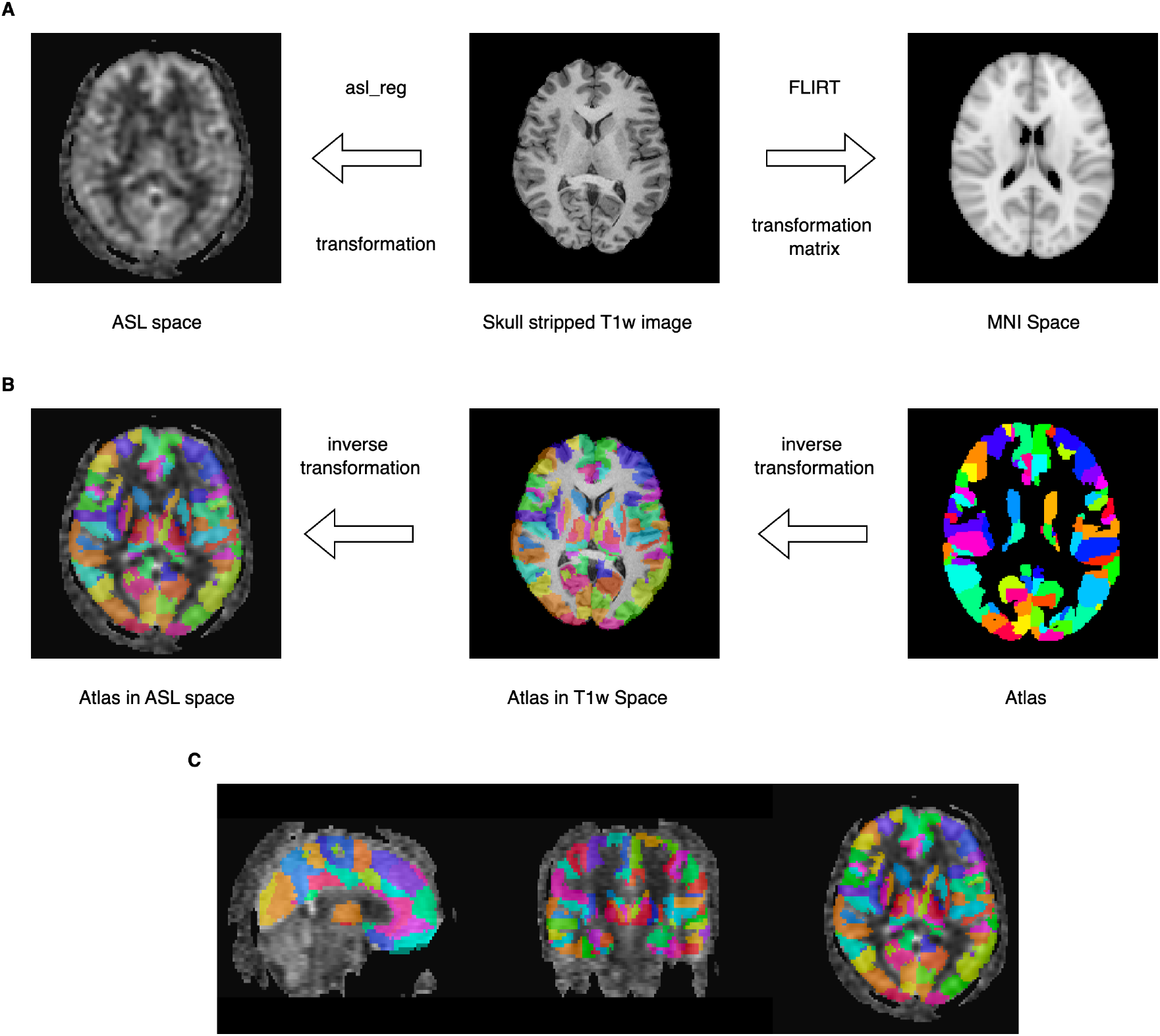
Multi-step registration pipeline for atlas-based cerebral blood flow quantification. (A) Forward transformation workflow: ASL images are registered to skull-stripped T1-weighted images using asl_reg from FSL’s Oxford ASL Toolbox to derive subject-specific T1w-to-ASL transformation matrices. Simultaneously, skull-stripped T1-weighted images are linearly registered to 1 mm MNI152 standard space using FLIRT with 12 degrees of freedom. (B) Inverse transformation for atlas mapping: the Brainnetome Atlas is transformed from MNI152 space to subject-specific T1-weighted space using the inverse transformation matrix computed with convert_xfm, then further transformed into ASL space using the T1w-to-ASL matrix. This multi-step approach ensures precise voxel-level alignment between the 246-region Brainnetome Atlas and low-resolution CBF maps while preserving spatial accuracy. (C) Final atlas alignment in ASL space shown in sagittal, coronal, and axial views, enabling accurate regional CBF quantification through voxel-level ROI matching. ASL = arterial spin labeling; FLIRT = FMRIB’s Linear Image Registration Tool; ROI = region of interest.

### 2.3 Machine Learning Classification Framework

Seven classifiers were evaluated using regional CBF features: Logistic Regression, Random Forest, XGBoost, Light-GBM, Support Vector Machine (linear and RBF kernels), and k-Nearest Neighbors. Features were standardized using z-scoring, and class imbalance (150 females, 95 males) was addressed via class weighting for all models except k-NN. Model performance was evaluated using stratified 5-fold cross-validation, with balanced accuracy as the primary metric. Additional metrics included ROC-AUC, accuracy, average precision, macro precision, macro recall, F1-score, and specificity. The same framework was applied to morphometric features for comparison.

A custom deep neural network (DNN) based on the Simple Fully Convolutional Network (SFCN) architecture was developed for voxel-wise classification using derived 3D CBF maps. The architecture comprised three 3D convolutional layers (64, 128, 64 filters, kernel size 3), max pooling, one 3D convolutional layer (kernel size 1), and global average pooling. The network contained ∼2.1 million trainable parameters. Hyperparameters were optimized via grid search over batch sizes (4–32), learning rates (10^*−*4^ to 5× 10^*−*3^), and optimizers (Adam, SGD). Weighted binary cross-entropy loss was used to address class imbalance. Training was performed using PyTorch 2.5.1 on NVIDIA A6000 GPUs. An 80/20 train-test split was used, followed by 5-fold cross-validation on the training set.

### 2.4 Explainable AI Analysis

Two complementary interpretability methods were used, matched to the granularity of each model class. SHapley Additive exPlanations (SHAP) was applied to the linear (regional CBF and morphometric) classifiers to obtain region-level biomarker rankings in atlas space, while Gradient-weighted Class Activation Mapping (Grad-CAM) was applied to the SFCN-CBF deep network to obtain voxel-level spatial attention maps. SHAP was deliberately not applied to the deep model because its high-dimensional voxel inputs lack a direct atlas-region correspondence, whereas the linear models operate natively on the 246 Brainnetome ROIs and admit closed-form Shapley values via the LinearExplainer.

For regional CBF and morphometric classifiers, SHAP’s LinearExplainer [25, 26] quantified feature importance. For morphometric models, SHAP values for volume, surface area, and thickness were aggregated by region. Statistical significance was assessed across 500 models (100 repetitions of 5-fold CV). ROIs ranked in the top 20% by SHAP values were counted, and significance was determined using binomial testing with Bonferroni correction. ROIs identified in more than 289 of 500 models (*p <* 0.05, corrected) were considered significant contributors.

To visualize voxel-level contributions from the 3D DNN models, Layer-wise Grad-CAM was applied using the Captum library. Importance maps were computed for each subject by targeting the 64-channel convolutional layer, then interpolated and normalized to generate group-level heatmaps for males and females, aiding interpretation of model focus regions.

### 2.5 Statistical Analysis

All statistical analyses were performed using R version 4.1.2, with significance set at *p <* 0.05. Between-sex differences in demographic variables (i.e., age, brain volumes) were assessed using two-sided independent *t*-tests. Machine learning model performance was evaluated using stratified five-fold cross-validation. Model stability was assessed using the coefficient of variation (CV) of balanced accuracy across folds. Regional CBF differences between males and females were assessed using Mann–Whitney *U* tests for regions identified as significant via SHAP analysis. Between-sex comparisons of mean cortical and subcortical CBF were performed using two-sample *t*-tests, and within-sex comparisons (cortical vs. subcortical) were tested using paired *t*-tests. Variance equality was assessed via *F* -tests, and Welch’s correction was applied where necessary. Finally, overlap between perfusion- and morphometry-based biomarkers was quantified using Jaccard similarity coefficients and Pearson correlation between binary significance vectors (246-dimensional, one per region).

## 3 Results

### 3.1 Participant Characteristics

A total of 215 participants (150 females, 95 males; age range: 18–30 years, mean age = 24.2± 3.1 years) were included after quality control. All individuals were right-handed Chinese Han participants from northern China. No significant sex difference was observed in age (two-sample *t*-test, *p* = 0.60).

Volumetric analyses revealed significant sex differences across most brain measures (Table 1). Males demonstrated significantly larger absolute volumes for total intracranial volume, total brain volume, gray and white matter compartments, and most subcortical structures (all *p <* 0.001). Only the accumbens area and cerebellar white matter showed no significant sex differences (*p >* 0.05).

### 3.2 Statistical Analysis of Cerebral Blood Flow Patterns

Statistical comparison of global (whole-brain, cortical, and subcortical) CBF values between males and females revealed significant sex differences across all aggregation methods (Table 2; Figure 3A). Females demonstrated significantly higher whole-brain CBF values: mean CBF (*t* = 9.29, *p* = 4.43 *×* 10^*−*16^, Cohen’s *d* = 1.28, large effect), median CBF (*t* = 9.30, *p* = 4.36 *×* 10^*−*16^, Cohen’s *d* = 1.28, large effect), and maximum CBF (*t* = 3.89, *p* = 1.36 *×* 10^*−*4^, Cohen’s *d* = 0.60, medium effect).

**Table 2.** **Statistical comparison of regional CBF values between males and females in whole-brain, cortical and subcortical ROIs** using two-sided *t*-tests. Welch’s correction applied where variance equality was rejected by *F* -test.

| Compartment | Aggregation | Test | $t$ | $p$-value | Mean diff. | Cohen’s $d$ | Effect |
| --- | --- | --- | --- | --- | --- | --- | --- |
| Whole brain | mean | Welch | 9.29 | $4.43 \times 10^{-16}$ | −10.27 | 1.28 | large |
| | median | Welch | 9.30 | $4.36 \times 10^{-16}$ | −10.40 | 1.28 | large |
| | maximum | Student | 3.89 | $1.36 \times 10^{-4}$ | −12.05 | 0.60 | medium |
| Cortical | mean | Welch | 9.41 | $2.40 \times 10^{-16}$ | −10.70 | 1.30 | large |
| | median | Welch | 9.32 | $4.33 \times 10^{-16}$ | −10.71 | 1.29 | large |
| | maximum | Student | 3.93 | $1.17 \times 10^{-4}$ | −12.15 | 0.60 | medium |
| Subcortical | mean | Welch | 6.12 | $9.16 \times 10^{-9}$ | −4.93 | 0.82 | large |
| | median | Welch | 6.07 | $1.20 \times 10^{-8}$ | −4.82 | 0.82 | large |
| | maximum | Student | 2.65 | $8.64 \times 10^{-3}$ | −7.97 | 0.41 | small |

**Figure 3.**
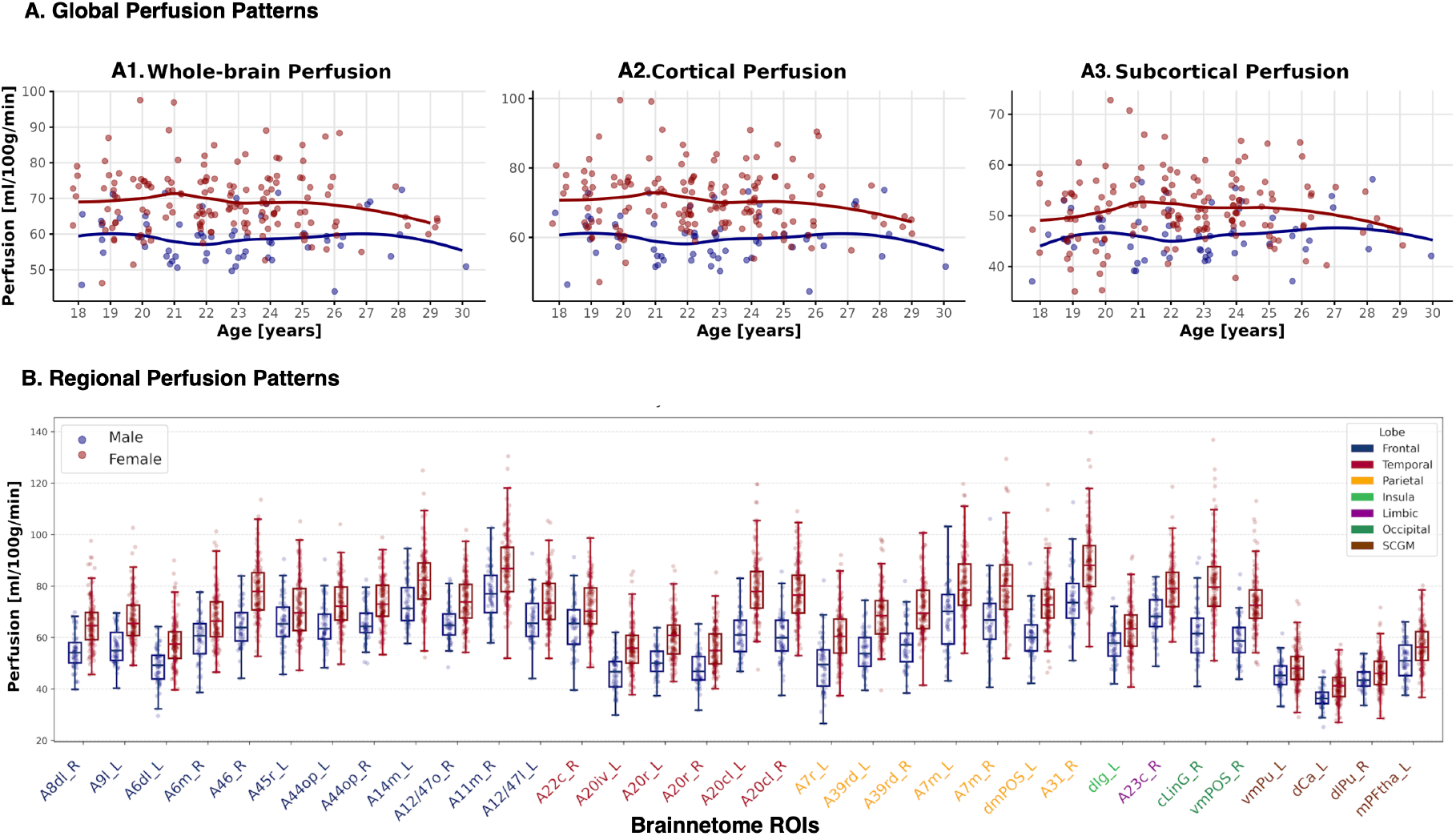
Multi-scale analysis of sex differences in cerebral perfusion. (A) Global perfusion patterns: scatter plots of mean CBF as a function of age for (A1) whole-brain, (A2) cortical, and (A3) subcortical compartments, with LOWESS smoothers fit per sex (males: blue; females: red). Females exhibit consistently higher CBF than males across all three compartments throughout the 18–30 year age range. (B) Local (regional) perfusion patterns at SHAP-identified discriminative regions: box plots of mean CBF in males (blue) and females (pink) across the 30 consensus sex-discriminative regions obtained from the intersection of mean- and median-CBF SHAP analyses (statistically significant at *p <* 0.05, Bonferroni-corrected; ranked in the top 20% of feature importance in *>* 289 of 500 logistic regression models). Region labels along the *x*-axis are colour-coded by lobe. CBF = cerebral blood flow; SHAP = SHapley Additive exPlanations; ROI = region of interest; SCGM = subcortical gray matter.

Regional analysis confirmed significant sex differences in both cortical and subcortical compartments. For cortical regions, females demonstrated significantly higher CBF values: mean CBF (*t* = 9.41, *p* = 2.40 ×10^*−*16^, Cohen’s *d* = 1.30, large effect), median CBF (*t* = 9.32, *p* = 4.33× 10^*−*16^, Cohen’s *d* = 1.29, large effect), and maximum CBF (*t* = 4.11, *p* = 1.17× 10^*−*4^, Cohen’s *d* = 0.60, medium effect). Similarly, subcortical regions showed consistent female superiority: mean CBF (*t* = 6.12, *p* = 9.16 ×10^*−*9^, Cohen’s *d* = 0.82, large effect), median CBF (*t* = 6.07, *p* = 1.20 ×10^*−*8^, Cohen’s *d* = 0.82, large effect), and maximum CBF (*t* = 2.86, *p* = 8.64 ×10^*−*3^, Cohen’s *d* = 0.41, small effect).

Within-sex analysis comparing cortical versus subcortical perfusion revealed that both males and females exhibited significantly higher cortical than subcortical CBF values. Both males and females showed large effect sizes across all measures (Cohen’s *d >* 2.0), indicating that the cortical-subcortical perfusion gradient is preserved across both sexes, with cortical regions consistently demonstrating higher perfusion than subcortical areas.

### 3.3 Data-Driven Classification Performance

Systematic evaluation of multiple machine learning models and feature representations revealed considerable variability in sex classification performance across imaging modalities (Table 3). The highest-performing model was a deep learning classifier trained on voxel-wise cerebral blood flow (CBF) maps using a modified Simple Fully Convolutional

**Table 3.**
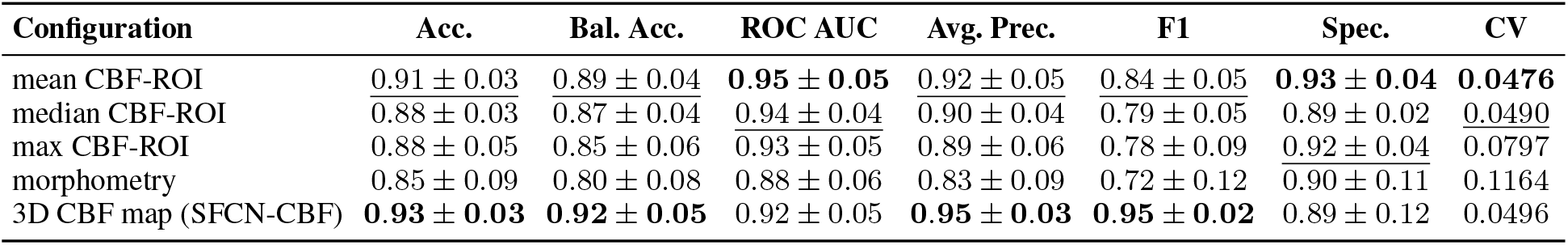
Sex Classification Performance Across Different Neuroimaging Modalities and Models. Performance metrics for sex classification using different neuroimaging features and machine learning approaches. Results are presented as mean ± standard deviation across 5-fold cross-validation. All ROI- and morphometry-based configurations use logistic regression; the 3D CBF map uses the SFCN-CBF deep network. Best result per metric in **bold**, second-best underlined. ROI = region of interest; Log Reg = logistic regression; SFCN = Simple Fully Convolutional Network; CBF = cerebral blood flow; Acc. = accuracy; Bal. Acc. = balanced accuracy; ROC AUC = receiver operating characteristic area under the curve; Avg. Prec. = average precision; Spec. = specificity; CV = coefficient of variance across folds. Macro-averaged recall is omitted as it equals balanced accuracy in the binary case.

Network (SFCN-CBF) (Figure 4), achieving a balanced accuracy of 0.92 ±0.05 and ROC-AUC of 0.92 ±0.05, with strong precision (average precision: 0.95 ±0.03) and F1 score (0.95 ±0.02). Among region-based CBF summaries, logistic regression using mean CBF values per ROI outperformed all other statistical summaries, achieving a balanced accuracy of 0.89 *±* 0.04 and ROC-AUC of 0.95 *±* 0.05. Median and maximum CBF values yielded slightly lower balanced accuracies (0.87 *±* 0.04 and 0.85 ± 0.06, respectively), though all variants maintained high specificity (median: 0.89 *±* 0.02, max: 0.92 *±* 0.03). Morphometric features derived from structural MRI offered substantially reduced discriminative power, with logistic regression yielding a balanced accuracy of 0.80 *±* 0.08 and ROC-AUC of 0.88 *±* 0.06.

**Figure 4.**
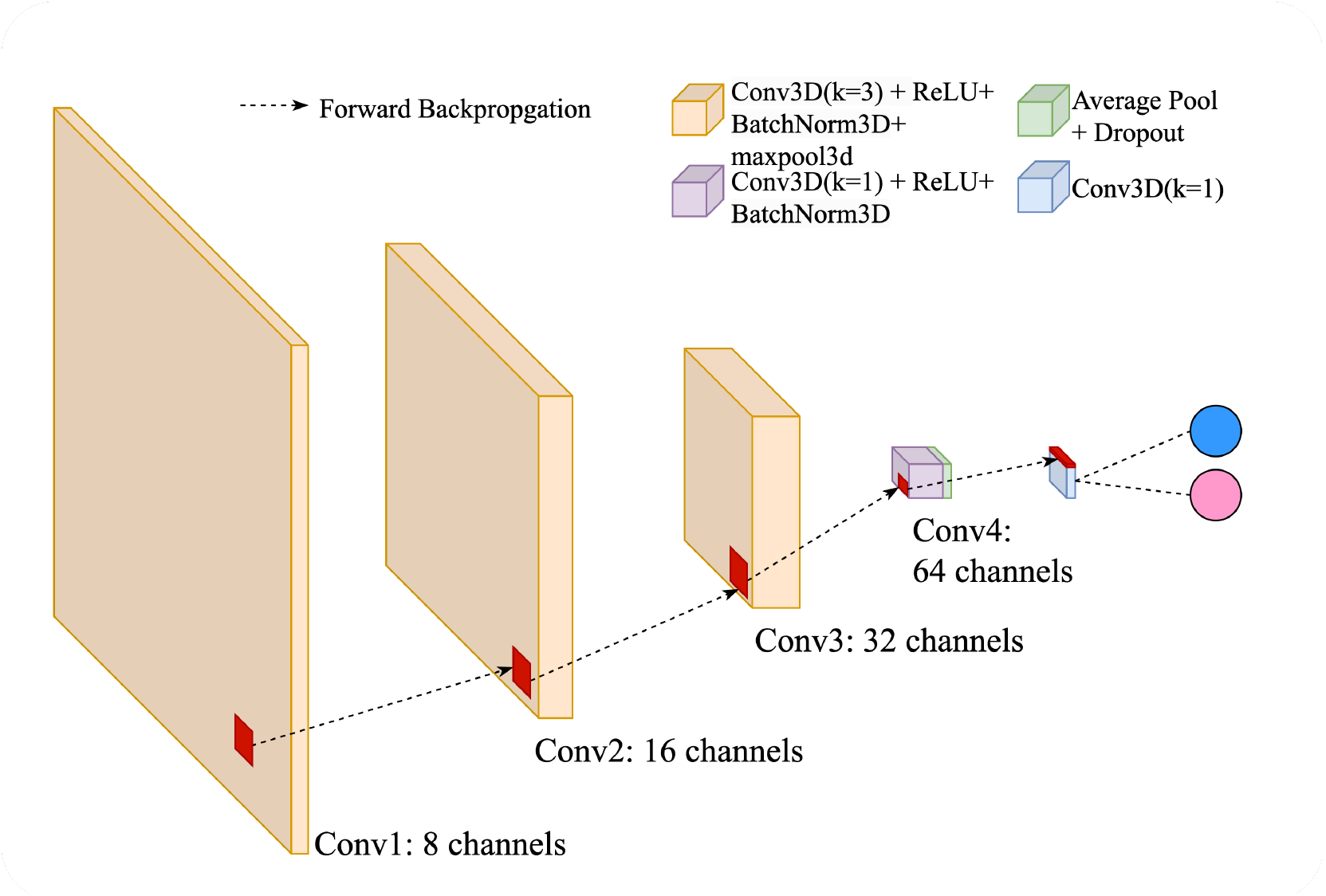
SFCN-CBF deep learning architecture. Modified Simple Fully Convolutional Network used for voxel-wise sex classification from 3D CBF maps. Three 3D convolutional layers (64, 128, 64 filters; kernel size 3) with max pooling, a 1*×*1*×*1 convolution, and global average pooling, totalling *∼*2.1 million trainable parameters.

These results demonstrate that CBF-based features provide superior discriminative power for biological sex classification compared to structural morphometry, with the deep learning approach on full CBF maps achieving the highest overall performance while regional CBF summaries offer competitive accuracy with greater interpretability.

To evaluate model stability across cross-validation folds, coefficient of variation (CV = std*/*mean) analysis revealed marked differences in model consistency. CBF-based models demonstrated superior stability, with mean CBF-ROI logistic regression exhibiting the highest stability (CV = 0.0476), followed closely by the 3D CBF deep learning model (CV = 0.0496) and median CBF-ROI model (CV = 0.0490). The maximum CBF-ROI model showed intermediate stability (CV = 0.0797), while morphometric features displayed substantially higher variability (CV = 0.1164). The consistently low CV values (*<* 0.08) across all CBF-based approaches indicate robust and reliable sex classification capabilities, whereas the higher variability observed in morphometric models suggests greater susceptibility to samplespecific variations and reduced generalizability. Given that the regional logistic regression model achieves accuracy within ∼ 3 percentage points of the deep network while operating natively on the 246 Brainnetome ROIs, we use it as the substrate for the SHAP-based biomarker analysis that follows, and reserve Grad-CAM for the SFCN-CBF model where voxel-level attention is the appropriate granularity.

### 3.4 SHAP-Based Identification of Discriminative Brain Regions

Explainable AI analysis using SHAP revealed robust sex-discriminative brain regions based on regional cerebral blood flow (Table 4; Figure 5). To identify aggregation-agnostic biomarkers, we examined the intersection between mean CBF (33 significant regions) and median CBF (37 significant regions) analyses, yielding 30 overlapping regions with high correlation (*r* = 0.84) and Jaccard similarity (*J* = 0.75). This strong concordance between mean and median CBF suggests minimal influence of outlier values and indicates consistent perfusion-based sex differences across statistical summaries.

**Table 4.** Distribution of sex-discriminative ROIs across lobes and networks. Counts of regions identified as significant sex-discriminative biomarkers by each modality and statistical aggregation. SCGM = subcortical gray matter; Mean *∩* Median = intersection of mean- and median-CBF biomarkers (consensus set).

| | Median | Mean | Max | Morph | Mean ∩ Median | Median ∩ Mean ∩ Morph |
| --- | --- | --- | --- | --- | --- | --- |
| *Hemisphere* |  |  |  |  |  |  |
| Left (L) | 19 | 17 | 18 | 15 | 16 | 2 |
| Right (R) | 18 | 16 | 22 | 13 | 14 | 2 |
| *Lobe* |  |  |  |  |  |  |
| Frontal | 13 | 12 | 14 | 3 | 11 | — |
| Parietal | 8 | 7 | 4 | 5 | 7 | 3 |
| Temporal | 7 | 6 | 11 | 8 | 5 | — |
| SCGM | 4 | 4 | 5 | 3 | 3 | — |
| Occipital | 3 | 2 | 2 | 6 | 2 | 1 |
| Limbic | 1 | 1 | 2 | 4 | 1 | — |
| Insula | 1 | 1 | 2 | 2 | 1 | — |
| *Network (Yeo 7)* |  |  |  |  |  |  |
| Frontoparietal | 10 | 8 | 7 | 3 | 8 | 2 |
| Default | 5 | 6 | 9 | 4 | 5 | — |
| Somatomotor | 5 | 3 | 8 | 2 | 3 | — |
| Visual | 5 | 3 | 4 | 9 | 3 | 1 |
| SCGM | 4 | 4 | 5 | 3 | 3 | — |
| Limbic | 3 | 4 | 5 | 2 | 3 | — |
| Ventral Attention | 3 | 3 | — | 2 | 3 | — |
| Dorsal Attention | 2 | 2 | 2 | 3 | 2 | 1 |
| **Total** | **37** | **33** | **40** | **28** | **30** | **4** |

**Figure 5.**
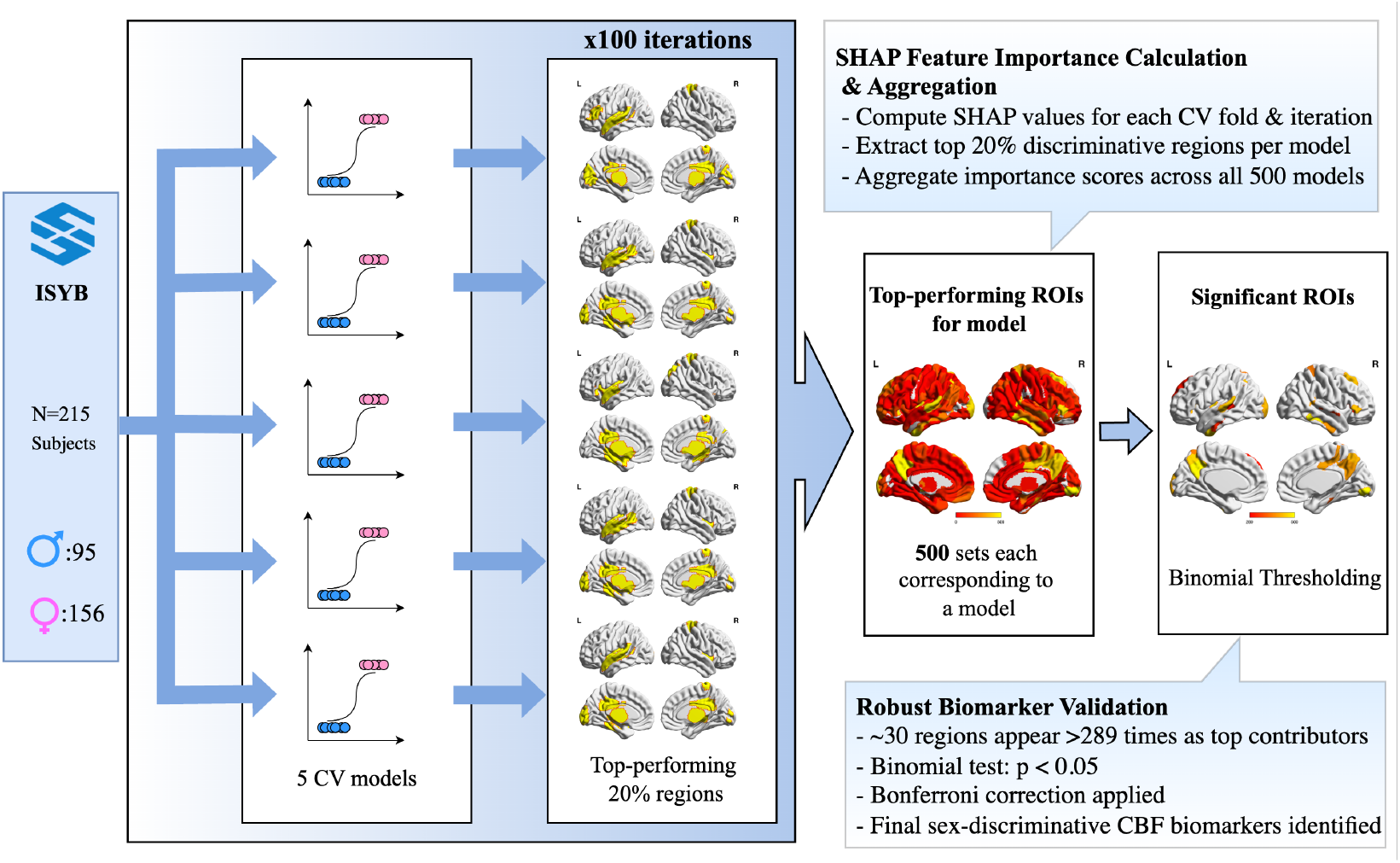
Explainable AI pipeline for identifying sex-discriminative cerebral perfusion biomarkers. SHAP-based model interpretation was performed across 100 iterations of 5-fold cross-validation on a sample of 215 subjects (male: 95; female: 150). For each of the 500 trained logistic regression models, SHAP values were computed to quantify the contribution of each brain region’s CBF to sex classification. Regions ranked in the top 20% of SHAP importance scores were identified per model. These top contributors were aggregated across all models, yielding a frequency map of regional importance. Statistical significance was determined via binomial testing, with Bonferroni correction applied for multiple comparisons. Regions appearing as top contributors in *>* 289 models (*p <* 0.05, corrected) were designated as significant sex-discriminative ROIs.

Region-level CBF distributions for males and females across these 30 consensus regions are shown in Figure 3B. The 30 consensus regions demonstrated pronounced anatomical clustering, with the frontal lobe showing the highest representation (*n* = 11, 37%), followed by the parietal lobe (*n* = 7, 23%) and temporal lobe (*n* = 5, 17%). Hemispheric distribution was nearly balanced (left: *n* = 16; right: *n* = 14). At the functional network level using Yeo 7-network parcellation, the frontoparietal control network showed the highest representation (*n* = 8, 27%), followed by the default mode network (*n* = 5, 17%). Independent Mann–Whitney *U* tests confirmed that all 30 regions exhibited statistically significant perfusion differences between males and females (all *p <* 0.05), validating that these SHAP-identified biomarkers reflect measurable biological variation.

SHAP analysis using maximum CBF values as regional summaries identified 40 statistically significant regions, representing the largest set among all CBF aggregation methods. However, these maximum-derived biomarkers showed reduced concordance with central tendency measures, exhibiting lower correlations and Jaccard similarities. Despite this reduced overlap with other CBF summary statistics, the maximum CBF model achieved competitive classification performance (balanced accuracy: 0.85 ±0.06), demonstrating that even extreme statistical representations can capture meaningful sex-discriminative information in cerebral perfusion patterns.

SHAP analysis of morphometric features identified 28 statistically significant regions with a markedly different anatomical and network profile compared to CBF-based biomarkers. The temporal lobe showed the highest representation (*n* = 8, 29%), followed by occipital (*n* = 6, 21%) and parietal lobes (*n* = 5, 18%). Network-level analysis revealed predominance of the visual network (*n* = 9, 32%) and default mode network (*n* = 4, 14%), contrasting sharply with the frontoparietal control network dominance observed in perfusion analysis.

### 3.5 Cross-Modal Biomarker Comparison

Cross-modal comparison between the 30 consensus CBF biomarkers and 28 morphometry-based discriminative regions revealed minimal overlap, with low Jaccard similarity (*J* = 0.074) and weak correlation (*r* = 0.023) (Figure 6). Only 4 regions showed significance in both modalities: A7r_L, A39rd_L, A39rd_R and cLinG_R representing 13.3% of CBF biomarkers. Notably, the A39rd region demonstrated bilateral representation across both hemispheres, suggesting robust structural-functional coupling in this parietal area for sex-related differences. Three of the four overlapping regions were in the parietal lobe, with network analysis showing predominant involvement of the frontoparietal control network.

**Figure 6.**
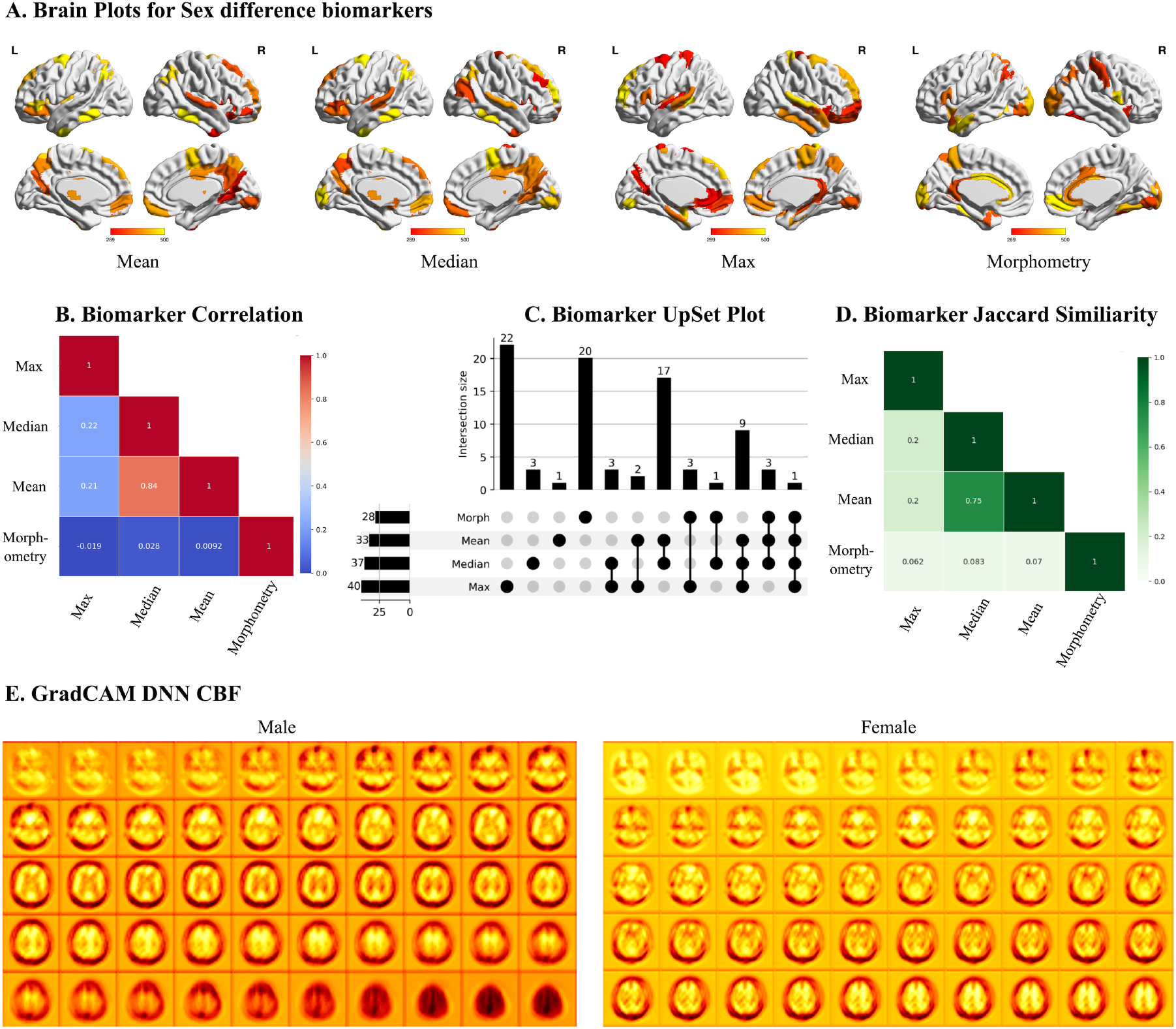
Multi-method explainable AI analysis of sex differences in cerebral perfusion. (A) Surface visualizations of brain regions identified as sex-discriminative biomarkers using mean, median, and maximum cerebral blood flow (CBF) measures, alongside morphometry-derived features. Warmer colors indicate stronger SHAP contributions. (B) Pairwise correlation matrix quantifying overlap in regions identified by each method. (C) UpSet plot showing intersections of biomarkers across modalities, highlighting both shared and unique contributions of mean, median, max CBF, and morphometry. (D) Radar plot summarizing the distribution of biomarkers across functional networks (Yeo 7-network atlas), demonstrating method-specific patterns. (E) Grad-CAM heatmaps from a deep neural network (SFCN-CBF model) trained for sex classification using CBF maps, visualized separately for males and females, capturing spatial regions of high model attention.

The consensus CBF biomarkers (mean ∩ median intersection) showed negligible correlation with morphometry (*r* = 0.022). The overall low cross-modal similarity (*J* = 0.074) indicates that cerebral perfusion and structural morphometry capture largely complementary aspects of sex-related brain differences, with perfusion measures providing superior discriminative power (balanced accuracy: CBF 0.89 *±* 0.04 vs. morphometry 0.80 *±* 0.08).

### 3.6 Deep Learning Model Interpretability

To analyse spatial contributions to sex classification, we applied Gradient-weighted Class Activation Mapping (GradCAM) to our SFCN-CBF model. Figure 6E displays group-level Grad-CAM activation maps averaged across male and female subjects. Both maps reveal predominant model attention to frontal lobe regions, consistent with our SHAP analysis identifying frontal areas as the most discriminative (37% of consensus biomarkers).

## 4 Discussion

The discriminative power of perfusion measures in distinguishing the sexes is leveraged using explainable data-driven approaches, pointing to a potential biomarker. It is also compared against traditionally used measures such as the morphometric features. The obtained findings and comparisons are described below.

### 4.1 Principal Findings and Clinical Implications

This study demonstrates that CBF implicitly carries superior discriminative power for biological sex classification compared to structural morphometry. This is evidenced by CBF-based models achieving 92% balanced accuracy (3D CBF map, SFCN-CBF), in comparison with morphometric features achieving only 80%. Notably, a simple logistic regression model on regional mean CBF achieved 89% balanced accuracy — within ∼3 percentage points of the ∼ 2-million-parameter deep network — pointing to the strength of the regional feature representation and the clinical viability of compact linear models that offer greater practicality and interpretability.

Because the linear and deep models converge to comparable accuracy, our explainable AI strategy was deliberately split: SHAP-based feature attribution was performed on the logistic regression models, where Shapley values admit closed-form computation under the LinearExplainer and yield region-level biomarker rankings that are directly interpretable in atlas space. Grad-CAM, on the other hand, was applied to the SFCN-CBF model to visualise voxel-level spatial attention from the deep network and to cross-validate the regional findings. This division leverages the best of both paradigms — accurate but opaque deep models for whole-image attention maps, and slightly less accurate but transparent linear models for ROI-level biomarker discovery — without sacrificing predictive performance, since the two modelling paradigms agree on the most discriminative anatomy (frontal-lobe dominance).

The study reports identification of 30 aggregation-agnostic ROIs (consistent with central tendencies, mean and median) that form perfusion biomarkers. These are predominantly localized in frontoparietal control networks, providing a neurobiologically interpretable framework for understanding sex differences in brain physiology. These findings suggest that perfusion measures are more sensitive to sex-related brain differences than traditional volumetric approaches, potentially offering enhanced utility for clinical applications requiring sex-stratified analyses.

### 4.2 Technical Advances in Perfusion Imaging for Sex Classification

#### ASL Perfusion Imaging: Optimal Modality Selection

The selection of arterial spin labelling over alternative perfusion modalities (SPECT, PET, DSC-MRI) provides critical advantages for sex classification. ASL’s non-invasive nature eliminates the necessity for toxic radioactive tracers or contrast agents, making it risk-free and enabling repeated measurements without safety concerns. ASL provides superior spatial resolution enabling precise anatomical localization within the 246-region Brainnetome Atlas. The quantitative nature yielding absolute CBF values (mL/100g/min) facilitates direct physiological interpretation, while integration with structural MRI provides comprehensive neuroanatomical context within a single session.

### 4.3 Neurobiological Mechanisms of Sex Differences in Cerebral Perfusion

The consistently higher range of CBF values observed in females across whole brain, cortical, subcortical and perfusion biomarkers (Cohen’s *d* = 0.82–1.30; large effect) reflect greater vascular coupling at all scales of evaluation. Estrogen’s vasodilatory effects on cerebral vasculature provide a primary hormonal explanation, with estradiol enhancing nitric oxide-mediated vasodilation and improving cerebrovascular reactivity. The young adult Chinese Han cohort considered here (18–30 years) captures these post-pubertal hormonal effects before age-related vascular changes, representing an optimal window for investigating sex-specific perfusion patterns.

The predominance of frontoparietal control network regions among discriminative biomarkers (27% of consensus regions) aligns with established sex differences in executive function, attention, and cognitive control. These networks demonstrate high metabolic demands and dense vascular supply, potentially amplifying sex-related perfusion differences.

Metabolic factors may also contribute, as females typically demonstrate higher glucose metabolism and different patterns of neural efficiency during cognitive tasks. The preservation of cortical-subcortical perfusion gradients across both sexes (Cohen’s *d >* 2.0) indicates that sex differences operate within conserved organizational principles of brain vascularization.

Notably, unlike morphometric measures, CBF is a quantitative intensity measure that is independent of intracranial volume; therefore total intracranial volume (TIV) does not act as a confound for CBF-based classification, unlike volume-matched structural classification approaches [22, 24].

### 4.4 Cross-Modal Biomarker Analysis and Clinical Translation

The minimal overlap between perfusion and morphometric biomarkers (Jaccard similarity = 0.074) demonstrates that these modalities capture complementary aspects of sex-related brain differences. The consensus approach identifying 30 robust biomarkers through mean–median CBF intersection provides a methodologically rigorous framework for biomarker discovery. The high correlation (*r* = 0.84) and Jaccard similarity (*J* = 0.75) between mean and median CBF analyses confirms minimal influence of outlier values, enhancing biomarker reliability for clinical translation. The bilateral representation of parietal regions (A39rd) across both perfusion and morphometric analyses suggests robust structural-functional coupling in areas critical for spatial processing and attention, where documented sex differences exist.

### 4.5 Limitations and Future Directions

Several limitations merit consideration. Our Chinese Han cohort, while providing ethnic homogeneity that reduces confounding, limits generalizability across populations with different genetic backgrounds and vascular risk profiles. The cross-sectional design cannot establish temporal stability of perfusion biomarkers or track developmental trajectories. ASL’s spatial resolution limitations compared to structural MRI may underestimate fine-grained perfusion differences, though our results suggest current resolution is sufficient for sex classification. Hormonal status (e.g., menstrual cycle phase) was not measured and may modulate cerebral perfusion. The SHAP LinearExplainer assumes feature independence, which may not strictly hold across spatially proximal ROIs. Replication in an independent, ethnically diverse cohort remains an important next step.

Future work should validate these biomarkers across diverse ethnic populations, examine temporal stability through longitudinal designs, and investigate integration with genetic and hormonal data to elucidate mechanistic pathways underlying sex differences in cerebral perfusion.

## 5 Conclusion

This study establishes cerebral blood flow as a superior biomarker for biological sex classification compared to structural morphometry, with simple linear models achieving performance comparable to complex deep learning approaches while offering enhanced clinical practicality. The identification of frontoparietal control network biomarkers provides neurobiologically interpretable targets for precision medicine applications, demonstrating the clinical utility of ASL perfusion imaging for sex-stratified brain analysis.

## Data and Code Availability

All analysis code is available at https://github.com/blackpearl006/superCBF (to be migrated to Zenodo). The ISYB dataset is available at https://www.scidb.cn/en/detail?dataSetId=826407529641672704. Region-wise CBF feature matrices for the 246 Brainnetome ROIs will be deposited at Zenodo upon publication.

## Author Contributions

N.A., N.S. and V.B. designed the research; N.A. and N.S. performed the research; N.A., N.S. and V.B. wrote the paper.

## Supplementary Figures and Tables

This section contains supplementary figures and tables that complement the main analyses. The mean-CBF biomarker boxplots are not duplicated here as they appear in the main text (Figure 3B).

**Figure S1.**
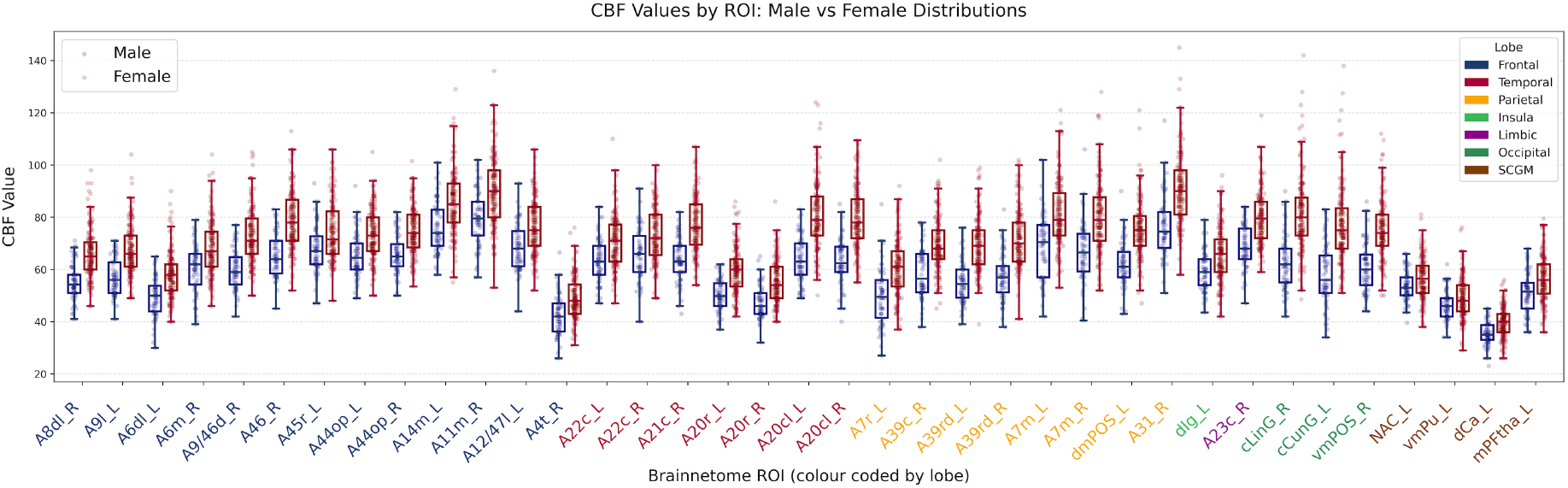
Sex-discriminative biomarkers identified using *median* regional CBF. Box plots of median CBF in males (blue) and females (pink) across the 37 SHAP-identified discriminative regions (*p <* 0.05, Bonferroni-corrected; ranked in the top 20% of feature importance in *>* 289 of 500 logistic regression models). Region labels are colour-coded by lobe. CBF = cerebral blood flow.

**Figure S2.**
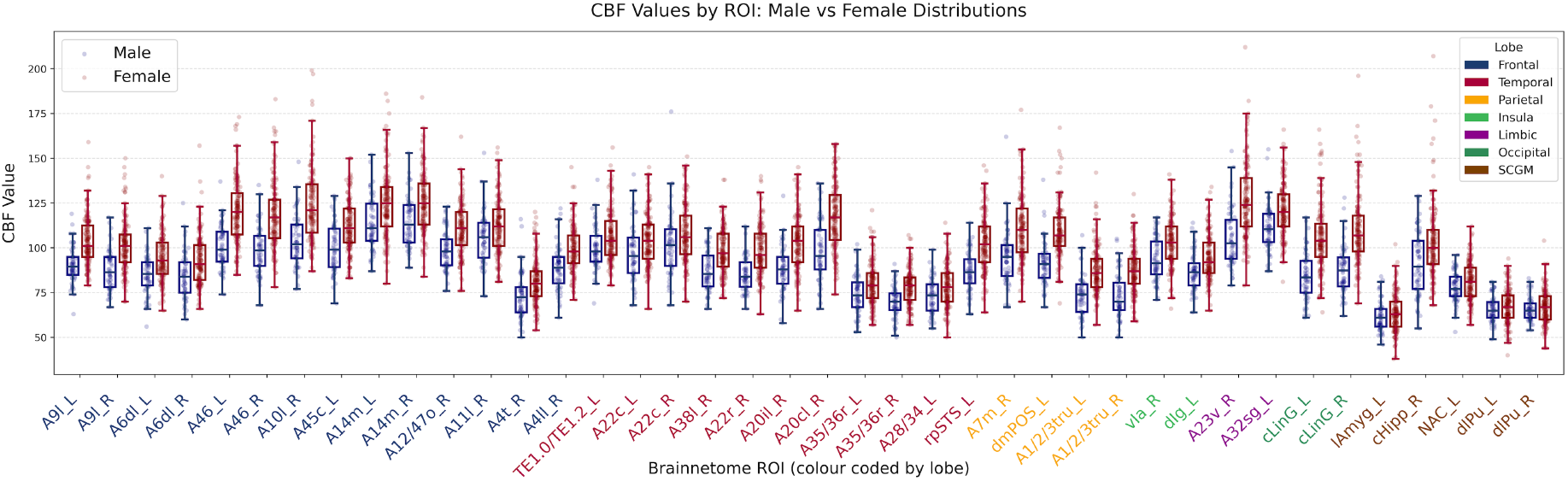
Sex-discriminative biomarkers identified using *maximum* regional CBF. Box plots of maximum CBF in males (blue) and females (pink) across the 40 SHAP-identified discriminative regions (*p <* 0.05, Bonferroni-corrected; ranked in the top 20% of feature importance in *>* 289 of 500 logistic regression models). Region labels are colour-coded by lobe. The maximum-CBF summary identifies the largest set of regions among all aggregation methods but with reduced concordance with mean and median analyses, as discussed in Section 3.4.

**Figure S3.**
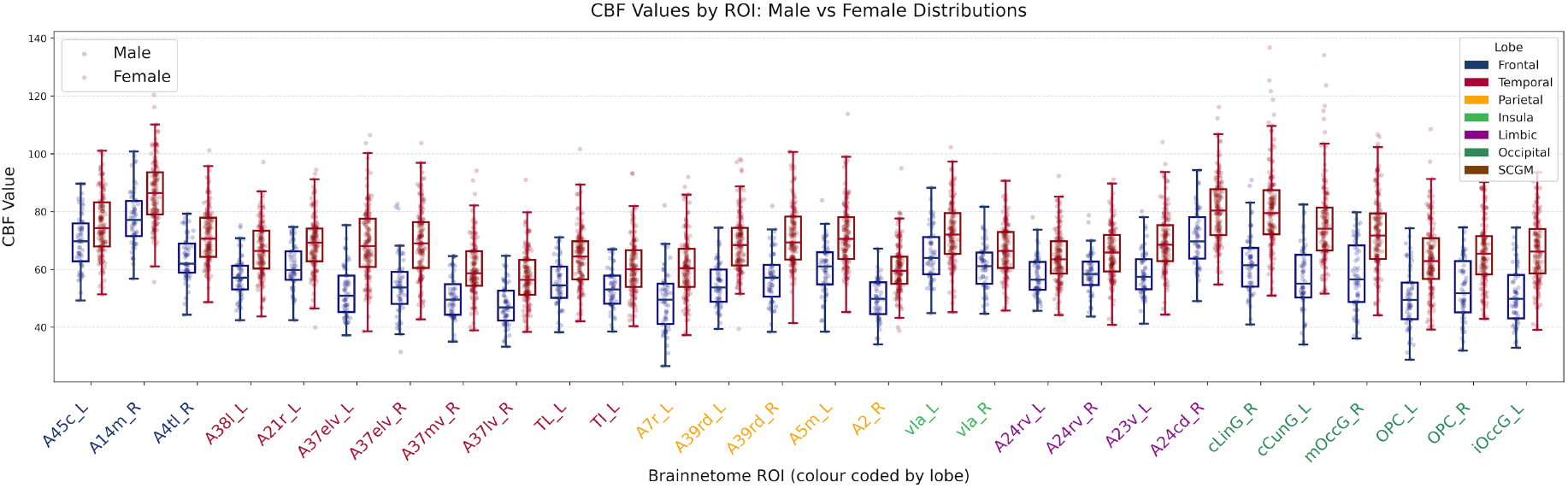
Sex-discriminative biomarkers identified from structural morphometry, visualised in the perfusion domain. Regions of interest are the 28 SHAP-identified morphometric biomarkers (computed from the regional aggregation of SHAP values across cortical thickness, surface area, and volume; *p <* 0.05, Bonferroni-corrected). For visualisation, *mean* CBF within each morphometry-identified ROI is plotted for males (blue) and females (pink) — i.e. the boxes show how much CBF differs between sexes inside regions selected on the basis of structural (not perfusion) features. The contrast with Figure 3B underscores the limited overlap (*J* = 0.074) between morphometry- and perfusion-derived biomarker sets reported in Section 3.5.

**Table S1.** Sex classification performance across all evaluated machine learning models and feature configurations. Mean± standard deviation across stratified 5-fold cross-validation. Best balanced accuracy per configuration in **bold**. Logistic regression and SVM (Linear) consistently outperform tree-based ensembles and k-NN across all CBF aggregations and morphometry, supporting the choice of logistic regression as the substrate for SHAP-based biomarker identification in the main analysis. Acc. = accuracy; Bal. Acc. = balanced accuracy; ROC AUC = receiver operating characteristic area under the curve; Avg. Prec. = average precision; Spec. = specificity. Small numerical differences from Table 3 reflect a separate model-sweep run with an independent random seed.

| Config. | Model | Acc. | Bal. Acc. | ROC AUC | Avg. Prec. | F1 | Spec. |
| --- | --- | --- | --- | --- | --- | --- | --- |
| mean CBF-ROI | Log Reg | 0.90 ± 0.07 | **0.87 ± 0.08** | 0.92 ± 0.05 | 0.91 ± 0.05 | 0.83 ± 0.12 | 0.94 ± 0.08 |
| | SVM (Linear) | 0.90 ± 0.03 | 0.86 ± 0.06 | 0.92 ± 0.05 | 0.91 ± 0.05 | 0.82 ± 0.06 | 0.95 ± 0.04 |
| | SVM (RBF) | 0.83 ± 0.06 | 0.80 ± 0.07 | 0.86 ± 0.06 | 0.83 ± 0.08 | 0.72 ± 0.10 | 0.89 ± 0.06 |
| | Random Forest | 0.84 ± 0.09 | 0.79 ± 0.11 | 0.83 ± 0.07 | 0.79 ± 0.08 | 0.71 ± 0.17 | 0.94 ± 0.08 |
| | XGBoost | 0.81 ± 0.09 | 0.77 ± 0.10 | 0.86 ± 0.04 | 0.82 ± 0.07 | 0.69 ± 0.15 | 0.88 ± 0.08 |
| | LightGBM | 0.81 ± 0.08 | 0.76 ± 0.09 | 0.84 ± 0.03 | 0.79 ± 0.06 | 0.68 ± 0.13 | 0.89 ± 0.08 |
| | k-NN | 0.83 ± 0.10 | 0.75 ± 0.14 | 0.83 ± 0.07 | 0.77 ± 0.12 | 0.64 ± 0.22 | 0.97 ± 0.06 |
| median CBF-ROI | Log Reg | 0.89 ± 0.05 | **0.86 ± 0.07** | 0.92 ± 0.06 | 0.91 ± 0.07 | 0.81 ± 0.09 | 0.92 ± 0.05 |
| | SVM (Linear) | 0.89 ± 0.04 | 0.85 ± 0.06 | 0.93 ± 0.06 | 0.92 ± 0.06 | 0.80 ± 0.07 | 0.94 ± 0.06 |
| | SVM (RBF) | 0.82 ± 0.07 | 0.78 ± 0.09 | 0.85 ± 0.07 | 0.82 ± 0.08 | 0.70 ± 0.13 | 0.89 ± 0.06 |
| | LightGBM | 0.82 ± 0.08 | 0.76 ± 0.11 | 0.85 ± 0.07 | 0.80 ± 0.08 | 0.67 ± 0.16 | 0.92 ± 0.05 |
| | XGBoost | 0.82 ± 0.05 | 0.76 ± 0.07 | 0.83 ± 0.05 | 0.80 ± 0.07 | 0.67 ± 0.10 | 0.92 ± 0.05 |
| | LightGBM | 0.80 ± 0.05 | 0.76 ± 0.07 | 0.83 ± 0.07 | 0.78 ± 0.08 | 0.66 ± 0.10 | 0.88 ± 0.04 |
| | k-NN | 0.81 ± 0.10 | 0.73 ± 0.13 | 0.82 ± 0.07 | 0.75 ± 0.12 | 0.61 ± 0.20 | 0.95 ± 0.06 |
| max CBF-ROI | Log Reg | 0.81 ± 0.07 | 0.76 ± 0.10 | 0.85 ± 0.05 | 0.82 ± 0.07 | 0.67 ± 0.14 | 0.89 ± 0.08 |
| | SVM (Linear) | 0.82 ± 0.05 | **0.78 ± 0.07** | 0.81 ± 0.06 | 0.79 ± 0.07 | 0.70 ± 0.10 | 0.89 ± 0.06 |
| | SVM (RBF) | 0.82 ± 0.05 | **0.78 ± 0.08** | 0.83 ± 0.05 | 0.79 ± 0.04 | 0.70 ± 0.11 | 0.89 ± 0.04 |
| | LightGBM | 0.80 ± 0.06 | 0.73 ± 0.09 | 0.81 ± 0.10 | 0.72 ± 0.14 | 0.61 ± 0.15 | 0.92 ± 0.05 |
| | XGBoost | 0.73 ± 0.12 | 0.66 ± 0.14 | 0.72 ± 0.10 | 0.63 ± 0.11 | 0.52 ± 0.21 | 0.85 ± 0.11 |
| | LightGBM | 0.71 ± 0.10 | 0.63 ± 0.11 | 0.72 ± 0.10 | 0.62 ± 0.12 | 0.47 ± 0.18 | 0.83 ± 0.12 |
| | k-NN | 0.75 ± 0.06 | 0.61 ± 0.09 | 0.77 ± 0.05 | 0.65 ± 0.09 | 0.34 ± 0.23 | 0.98 ± 0.03 |
| morphometry | Log Reg | 0.85 ± 0.08 | **0.80 ± 0.08** | 0.88 ± 0.06 | 0.83 ± 0.09 | 0.72 ± 0.12 | 0.90 ± 0.10 |
| | SVM (Linear) | 0.85 ± 0.06 | 0.79 ± 0.08 | 0.87 ± 0.06 | 0.82 ± 0.09 | 0.70 ± 0.11 | 0.92 ± 0.06 |
| | LightGBM | 0.83 ± 0.08 | 0.77 ± 0.08 | 0.86 ± 0.08 | 0.78 ± 0.08 | 0.68 ± 0.10 | 0.89 ± 0.11 |
| | XGBoost | 0.82 ± 0.08 | 0.76 ± 0.07 | 0.84 ± 0.07 | 0.78 ± 0.08 | 0.67 ± 0.11 | 0.90 ± 0.09 |
| | SVM (RBF) | 0.82 ± 0.09 | 0.76 ± 0.10 | 0.84 ± 0.08 | 0.78 ± 0.11 | 0.66 ± 0.15 | 0.90 ± 0.10 |
| | Random Forest | 0.80 ± 0.05 | 0.68 ± 0.05 | 0.84 ± 0.07 | 0.76 ± 0.08 | 0.53 ± 0.10 | 0.95 ± 0.07 |
| | k-NN | 0.79 ± 0.05 | 0.64 ± 0.07 | 0.72 ± 0.09 | 0.54 ± 0.13 | 0.44 ± 0.15 | 0.97 ± 0.04 |

